# The SpliZ generalizes “Percent Spliced In” to reveal regulated splicing at single-cell resolution

**DOI:** 10.1101/2020.11.10.377572

**Authors:** Julia Eve Olivieri, Roozbeh Dehghannasiri, Julia Salzman

## Abstract

To date, detecting robust single-cell-regulated splicing is viewed as out of reach from droplet based technologies such as 10x Chromium. This prevents the discovery of single-cell-regulated splicing in rare cell types or those that are difficult or impossible to sequence deeply. Here, we introduce a novel, robust, and computationally efficient set of statistics, the Splicing Z Score (SpliZ) and SpliZVD, to detect regulated splicing in single cell RNA-seq including 10x Chromium. The SpliZ(VD) provides annotation-free detection of differentially regulated, complex alternative splicing events. The SpliZ generalizes and increases statistical power compared to the Percent Spliced In (PSI) and mathematically reduces to PSI for simple exon-skipping. We applied the SpliZ to primary human lung cells to discover hundreds of genes with new regulated cell-type-specific splicing. The SpliZ has wide application to enable biological discovery of genes predicted to have functionally significant splicing programs including those regulated in development.

## Main Text

Splicing is a core function of eukaryotic genes that generates proteins with diverse and even opposite functions^1^, changes translation efficiency^2^, controls localization^3^, and generates non-coding RNAs^4^. Very few genes’ splicing programs have been characterized at single-cell resolution, and the function of splicing remains a critical open problem in biology^5^. Constant advances in single-cell RNA sequencing (scRNA-seq) technology now provide an unprecedented opportunity to understand how splicing is regulated at the single cell level.

The enormous complexity of splicing in eukaryotic genomes and the low sequencing depth per cell in scRNA-seq experiments makes it challenging to precisely quantify RNA isoform expression and its differential regulation in single cells. With few exceptions^6^, the field has typically attempted to either estimate isoform expression using model-based approaches^7–10^ and then perform differential splicing analysis, or directly quantify exon inclusion^7,8^ using percent spliced-in (PSI).

Each such approach has significant drawbacks, especially in scRNA-seq. First, annotation-based methods for isoform quantification give notoriously unstable point estimates in the presence of non-uniform read sampling^9^ or when annotations are incomplete or erroneous. These problems are exacerbated at low sequencing depths. Further, droplet-based scRNA-seq methods are 3’-biased, making accurate isoform estimation impossible even with deep sequencing.

The second approach is to use PSI, which quantifies the fraction of transcripts that skip a specific exon^11^. However, tests based on PSI must proceed exon-by-exon, requiring hundreds of thousands of tests, and cannot detect splicing events beyond simple exon skipping. This diminishes statistical power through multiple hypothesis testing, and creates problems with statistical dependency when testing differential exon skipping multiple times per gene. Moreover, PSI is thought to be statistically unreliable for capturing exon skipping with less than 10 recovered mRNA on average per exon skipping event^11^.

For these reasons, published methods for analyzing splicing in scRNA-seq are almost exclusively inappropriate for use on droplet-based data^12,13^. Those that are recommended for droplet-based analysis rely on genome annotation, require pairwise tests between all cell types which scale poorly for many cell types and lose sensitivity due to multiple hypothesis testing, and only analyze exon-skipping^14^. These issues have led to the view that robust differential splicing analysis in droplet-based scRNA-seq is out of reach^11,15,16^.

Here we introduce the Splicing Z Score (SpliZ), a scalar value assigned to each gene-cell pair that quantifies how deviant a cell’s splicing is compared to a population average. The SpliZ can be applied to compiled junctional read counts from any single cell data (e.g. STAR-aligned^17^ and SICILIAN^17,18^-processed plate or droplet-based scRNA-seq data) and, even more generally, to bulk data, though that is not the focus of this work. The SpliZ integrates all non-constitutive spliced reads on a per-gene basis to detect deviant splicing patterns in single cells under low and biased sampling (Fig. 1A). It also requires one test per gene, greatly reducing the number of tests from potentially 10s (or in some cases, 100s) required by PSI. As shown below, the SpliZ has power to detect isoform expression changes beyond exon skipping.

**Figure 1.**
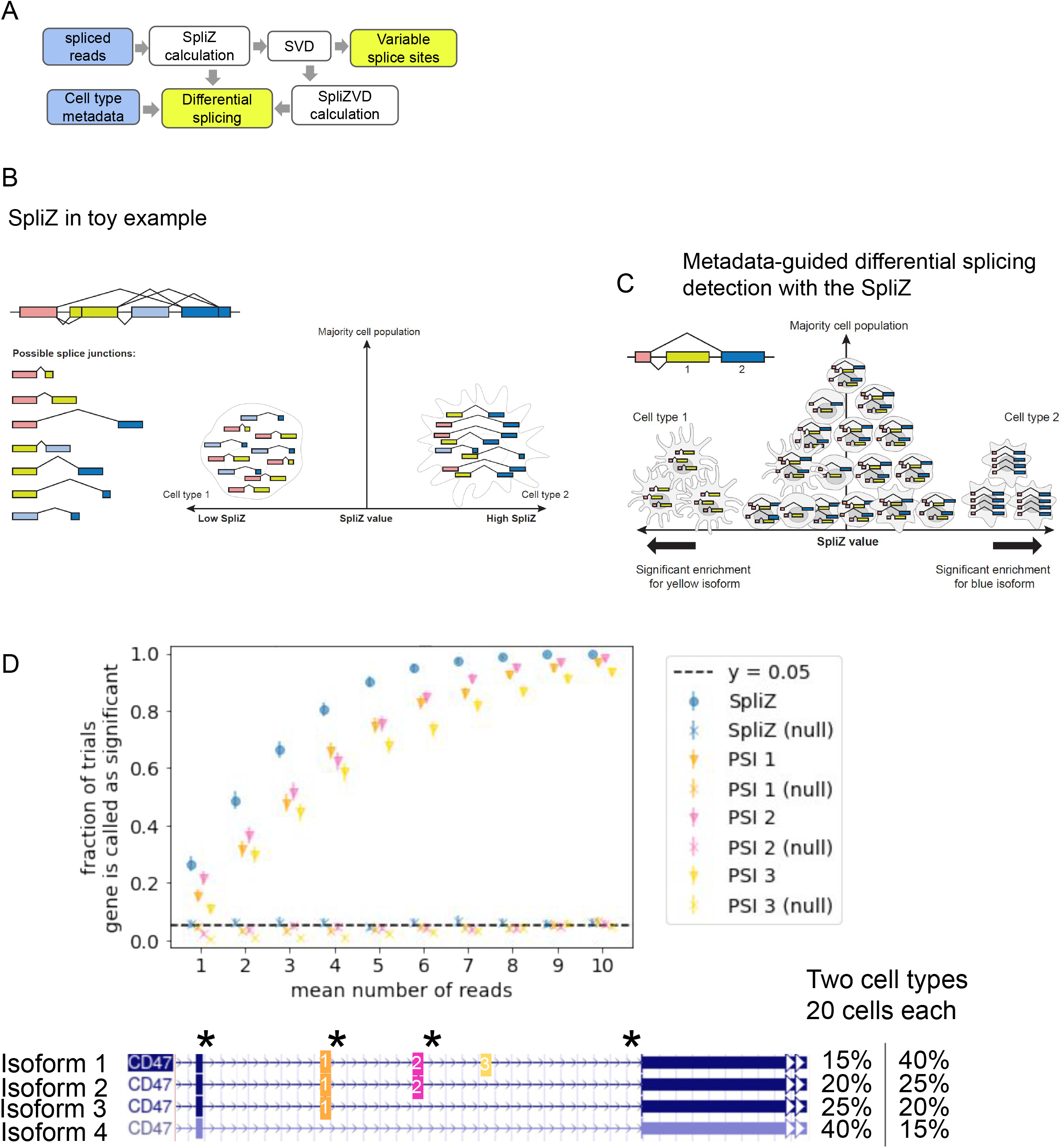
(A) The SpliZ takes spliced reads and metadata and returns genes called as differentially spliced by cell type. (B) SpliZ toy example: cell on the left has short average introns vs the cell on the right, giving it a lower SpliZ. (C) SpliZ scores can be aggregated for each cell type, identifying cell types with statistically deviant splicing. (L) cell type with significant enrichment for short introns; (R) cell type with significant enrichment for long introns. (D) Simulations of the SpliZ, SpliZVD, and PSI: In both, two cell types with 20 cells respectively are simulated, each having a different proportion of isoforms (1000 trials). At each read depth, Poisson(n) reads are sampled in proportion to isoform abundances. Null values are calculated from cell populations with identical isoform expression. Driving splice sites are starred by asterisks and coincide with simulated alternative splice sites. The SpliZ increases power over PSI for cell types expressing different proportions of CD47 isoform structure.

The SpliZ quantifies cell-type-specific splicing for each cell-gene pair by (1) assigning each read aligning to a splice junction to a rank based on the relative size of the intron compared to the set of observed introns for that 3’ and 5’ splice site, respectively (Methods); (2) converting the rank to a residual measuring its statistical deviation compared to “the typical” intron length rank (Methods); (3) statistically grounded scaling and summing of these values to a single SpliZ per gene and cell. A cell-gene pair has a large negative (resp. positive) SpliZ, if, on average, introns have statistically significantly smaller (resp. longer) intron length compared to the population of cells profiled in an experiment (Fig. 1B).

The SpliZ is a statistic developed from first principles and has the important property that it is a formal generalization of PSI (Methods), mathematically reducing to PSI in the case of exon-skipping with only two measured junctions. The SpliZ increases power over PSI in cases of more complicated splicing. If there are multiple differential exon skipping events in the same gene, PSI lacks power to detect each difference, while the SpliZ builds strength across both events, resulting in increased power.

The theoretical properties of the SpliZ allow it to be efficiently integrated into significance testing: Under the null hypothesis, the SpliZ has mean 0 for every cell type, while under the alternative hypothesis each cell type has a constant mean SpliZ value, not all of which equal zero (Fig. 1C, Methods). A common bioinformatics practice approximates p values, in this case to test for differential splicing by cell type in each gene, using permutation distributions. However, this approach, while intuitive, has been proven to dramatically underestimate the type I error in the case of unequal variance between cell types^19^. Further, permutation testing is computationally intensive, requiring (number of genes)*(number of permutations) evaluations. To overcome this computational and statistical problem, we adopted a procedure with better type I error control^19^ and developed a two-step method to estimate significance: First, the SpliZ is referred to a theoretical null distribution. For p-values passing a nominal 0.05 level, permutations are performed to estimate the p value with higher precision. This results in (a) a conservative FDR estimate (b) 95% less computational cost under a global null distribution. Final p values are adjusted using the Benjamini-Hochberg correction^20^. By performing only one test per gene, and determining differential alternative splicing between cell types by one gene-wide test rather than pairwise tests between cell types, this method avoids reduction in statistical power by removing unnecessary multiple hypothesis testing.

As predicted by theory, the SpliZ has higher power than PSI when multiple exons are skipped in the same transcript. When there are multiple isoforms for a gene, each of which includes one more exon, the SpliZ builds strength across these isoforms to identify real differences between cell types at lower read depths than PSI, while maintaining the same type II error rate (Fig. 1D). Although all spliced reads are used both to compute both the SpliZ and PSI, three different PSI values must be computed for each non-constitutive exon, yielding three scores each of which have less power than the SpliZ to detect differences in isoform usage across the two cell populations. At an average of five reads per cell, there is a 90% chance of a difference being detected by the SpliZ, but less than an 80% chance of a difference being detected by PSI.

While the SpliZ has high statistical power to detect exon skipping and other complex patterns (Fig. 1D), it lacks power to detect isoform shifts in some cases, for example between isoforms with alternative cassette exons. To overcome this lack of power, we developed the SpliZVD, a modification of the SpliZ, which as suggested by the name, is a gene-cell score computed by projecting splicing residuals onto eigenvectors of its SVD (we consider only the first projection in this work) (Fig. 2A, Methods). The SVD of the residual matrix used to compute the SpliZVD is also used for biological interpretability of the SpliZ and SpliZVD. Splice sites corresponding to up to the three largest-magnitude components of the first eigenvector are nominated as the statistically most variable splice sites (Fig. 2B, Methods).

**Figure 2.**
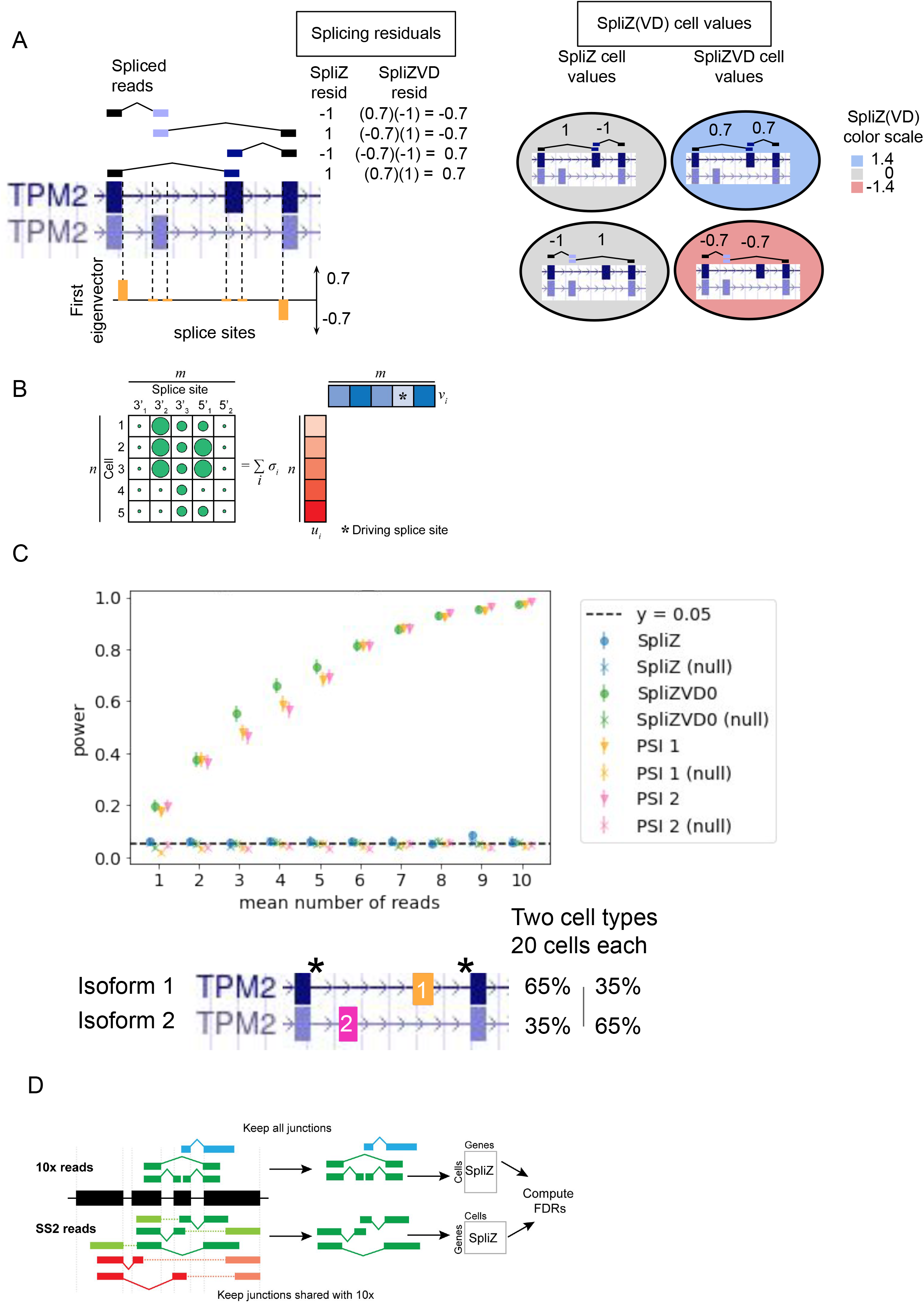
(A) The SpliZVD is the projection of the matrix of splicing residuals onto its first eigenvector. (B) The SVD is used to identify the most variably alternatively spliced sites. (C) Simulation as described in 1D. As predicted by theory, the SpliZVD calls differential alternative splicing between the two cell types with different proportions of TPM2-like isoforms, while the SpliZ does not. (D) The SpliZ is calculated independently for Smart-seq2 data restricted to junctions detected by 10x to measure technology-dependence of results.

The SpliZVD maintains the power of PSI in simulation with two cassette exons, a situation where the SpliZ has no power to detect differences because both isoforms contain one “long” and one “short” intron (Fig. 2C). In both this and the previous simulation, the splice sites involved in differential alternative splicing were correctly identified by the method (Fig. 1D, Fig 2C). The SpliZ and SpliZVD together have been shown both through theory and simulation to identify splicing differences with as much power as PSI or more, while extending to more complicated splicing patterns.

We analyzed 60,550 carefully annotated human lung cells from the Human Lung Cell Atlas (HLCA)^21^ from two individuals to test the performance of the SpliZ and SpliZVD on real scRNA-seq data. The HLCA, sequenced on the 10x (53,469 cells) and Smart-seq2 (7,081 cells) platforms, includes 57 annotated cell types with highly variable depth of sampling per cell type. We focused the SpliZ analysis on 10x because orders of magnitude more cells are profiled through the technology, including cell types that cannot be profiled or are poorly sampled by Smart-seq2, and ran the pipeline on Smart-seq2 restricted to junctions found in the 10x data for comparison and validation purposes (Fig. 2D, Supplement). The SpliZ pipeline took under 3 hours and 80 GB to run on all analyzed data in parallel. A median of 912 genes are detected per cell in 10x data (1,660 in Smart-seq2 data), and a median of 67 genes have a computable SpliZ per cell for 10x data (833 for Smart-seq2 data). 1,754 genes have a computable SpliZ in at least 10 cells in one of the individuals (11,640 for Smart-seq2 data) (Methods).

The SpliZ and SpliZVD identified hundreds of differential alternative splicing events between cell types in 10x data from human lung. 210 (resp. 219) genes were called as having significant cell-type-specific splicing by the SpliZ, and 133 (141) genes were called by the SpliZVD, with 89 (90) genes called by both (p < 0.05). 178 genes were called by either the SpliZ or SpliZVD in both individuals’ 10x data (p value: 1.11e-16). There is significant correlation between median SpliZ scores for the same gene and cell type for 10x and Smart-seq2 data in the same individual, restricting to genes that are significant in both (Pearson correlation of 0.315 and 0.650).

Genes with the most deviant SpliZ values (Methods) include ATP5F1C, a core component of the mitochondrial ATP-synthase machinery, and MYL6, an essential component of the actin cytoskeleton with partially characterized splicing, both of which we find have cell-type-specific exon skipping events (Supp. Fig. 1A-B, Supp. Table 1, in preparation). Another of the many examples of highly cell-type-specifically spliced genes discovered is LMO7, an emerin-binding protein with alternative splicing affecting a protein domain of unknown function (Fig 3A).

**Figure 3.**
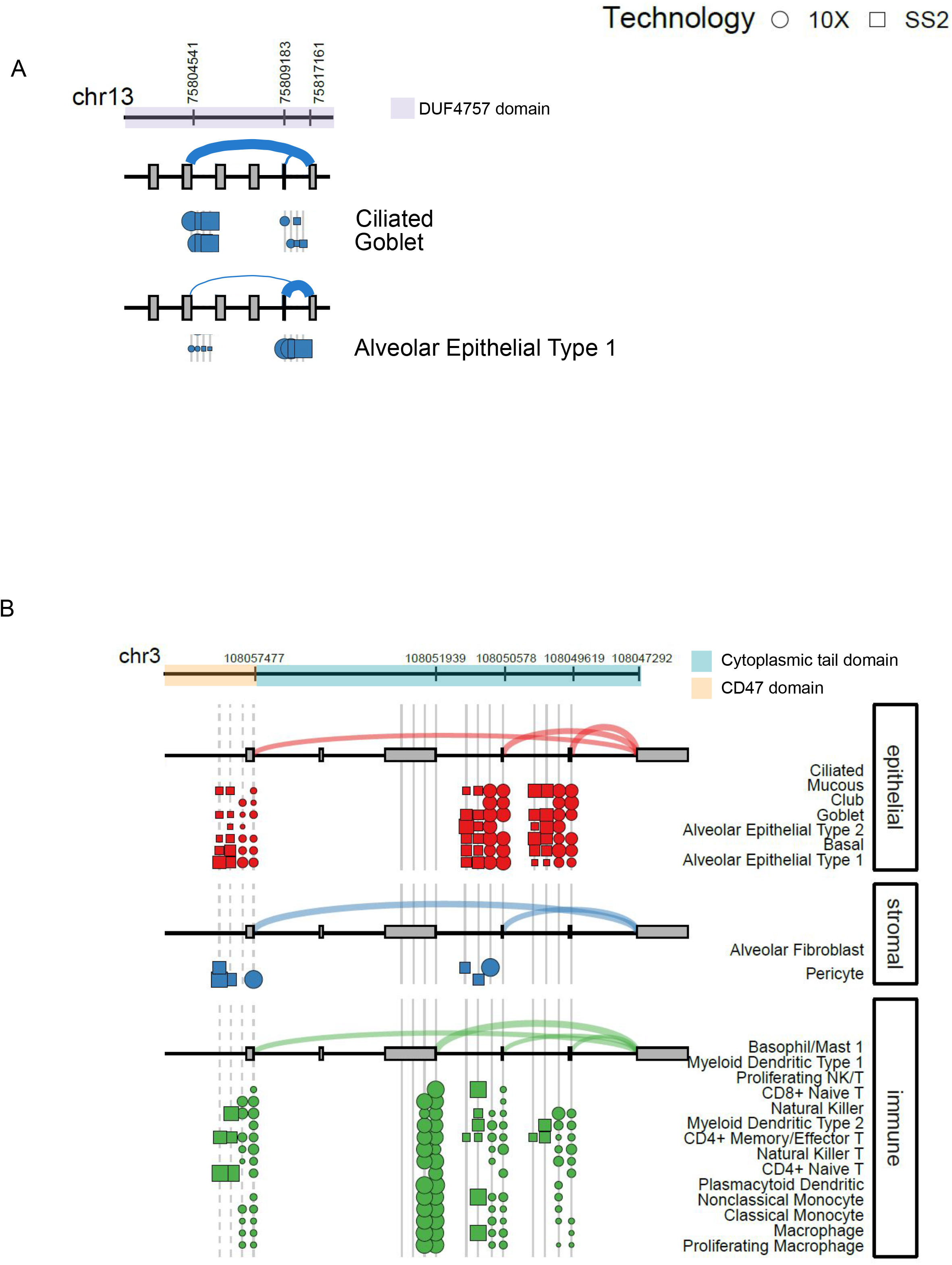
(A) LMO7 has significant splicing differences within epithelial cell types at splice site 75817161. Dot size represents the fraction of reads from each 5’ splice site to the 3’ site at 108047292. The arcs show the average site use across the tissue compartment,thicker arcs corresponding to higher fractional use. Read fractions from two 10x and Smart-seq2 datasets are shown by dots and squares, respectively. 92% of reads skip the proximal exons, compared to 2% in alveolar epithelial type 1 cells in Smart-seq2 data. (B) Differential splicing in CD47 is tissue-compartment-specific. 6% of reads from 10x splice to an unannotated 5’ splice site (dashed lines).

The SpliZ identifies a pattern of differential alternative splicing involving at least three isoforms in CD47 (Fig. 3B, Supp. Table 1). CD47 is an immune-regulatory membrane protein that has recently been identified as a therapeutic target in a set of myeloid malignancies^22^ but has undescribed cell-type-specific splicing programs. Automatic detection of the most variable splice sites for CD47 identified the 5’ and 3’ splice sites flanking a region with three exons that can be sequentially included to create four different isoforms, just as in Figure 1D. The SpliZ builds strength across these multiple skipping events to identify differential alternative splicing in the gene, with cells in the epithelial compartment tending to express three different isoforms equally while the other compartments are dominated by just one of the isoforms.

In real biological scenarios the SpliZVD improves power to detect differential splicing for some genes: 44 (51) genes are called by the SpliZVD but not the SpliZ in individual 1 (2). Although the tropomyosin gene TPM2 is called by both the SpliZ and SpliZVD, it only had the 266 (204)th-largest-magnitude maximum ontology median by the SpliZ, it had the 121 (82) largest by the SpliZVD in individual 1 (2). This differential alternative splicing event is also called as significant by Smart-seq2 in the second individual with the SpliZVD. Increased power of the SpliZVD over the SpliZ would be predicted due to the cassette exon gene structure of TPM2, which is analogous to the gene structure in Figure 2C. The residual matrix’s SVD automatically identifies the most variable splice sites for TPM2: a 3’ splice site bordering the two cassette exons, as well as a 5’ splice site that can splice to two different transcript ends.

Overall, ~28% (147 out of 529) of the most variable splice sites for significantly differentially spliced genes (p<0.05) in the HLCA corpus are annotated alternative exons (Methods). One of the genes with the highest magnitude median SpliZ is PPP1R12A, a protein phosphatase regulatory subunit: its most variable splice site is not annotated as being alternatively spliced (Supp. Fig. 1C). PPP1R12A is also called as by the SpliZVD in Smart-seq2. This supports the use of the SpliZ to re-identify known alternative splicing events and as a purely statistical, annotation-free, approach to discover new alternative splicing events that are cell type-specifically regulated.

Undoubtedly, the 3’ bias of 10x genomics data derived from priming on the poly-A tail restricts discovery of regulated splicing in this study by biasing data and measurement towards only a subset of splice sites. Moreover, because the SpliZ collapses all splice events to a single scalar, it lacks power to detect some differential splicing events. The SpliZVD remedies this problem through a data-driven approach based on variance-maximizing projections of splicing residuals which are especially important for statistical power when transcripts are profiled more deeply and more uniformly and isoform structure is complex. Indeed, as predicted by theory, the SpliZVD detects significantly more events than the SpliZ in Smartseq-2, explored in other work (in preparation). Also note that the methodology in the SpliZ pipeline can be applied to any RNA-seq dataset, including bulk sequencing.

Because the SpliZ has a tractable statistical distribution and is single-cell resolved, it enables new biological analysis beyond the scope of this paper, including clustering approaches on the basis of the SpliZ alone or in combination with gene expression. In addition, the SpliZ can be correlated with any other numerical phenotype, such as pseudotime, cell cycle trajectory, gene expression or spatial position in the context of single cell spatial genomics. In summary, by deconvolving technical and biological noise in splicing, the SpliZ provides a new method for the field to identify and prioritize splicing events that are regulated and functional and to address decades-long debates regarding the degree to which alternative splicing is regulated at a single cell level.

## Acknowledgments

We thank Peter Wang and Steve Quake for insight and instrumental comments during the development of the method, and Elisabeth Meyer and Robert Bierman for comments on the manuscript. Thank you to Jessica Klein for creating parts of figure 1 and 2, and also Emma Chory for figure assistance. We thank Kyle Travaglini and Mark Krasnow for providing access to the HLCA data prior to publication. J.O. is supported by the National Science Foundation Graduate Research Fellowship under Grant No. DGE-1656518 and a Stanford Graduate Fellowship. R.D. is supported by the Cancer Systems Biology Scholars Program Grant R25 CA180993 and Clinical Data Science Fellowship Grant T15 LM7033-36. J.S. is supported by the National Institute of General Medical Sciences Grant R01 GM116847 and NSF Faculty Early Career Development Program Award MCB1552196.

## Methods

### 1 Code Availability

The SpliZ code along with the codes used for data analysis and to create the figures are available through a GitHub repository: https://github.com/juliaolivieri/SZS_pipeline/. Installed package versions (also available in environment.yml file on github): matplotlib [6]: 2.2.3; numpy [5]: 1.18.4; pandas [7, 12]: 1.0.4; pickle [10]: 4.0, scipy [11]: 1.4.1; statsmodels [8]: 0.11.1; tqdm [2]: 4.46.0.

### 2 Data Availability

Human lung cell atlas (HLCA) data was downloaded from the EGA archive at accession number EGAS00001004344 [9]. We refer to Patient 2 in HLCA as Individual 1 and Patient 3 as Individual 2 in this manuscript. The cell types we use here are based on concatenating the “compartment” and “free annotation” columns from the HLCA metadata and only considering lung cells (not blood).

#### Preparing splice junction input files

The scRNA-Seq data sets were mapped to the reference human genome (GRCh38) using STAR version 2.7.5a with default parameters [4].

We used SICILIAN [3] for calling splice junctions. SICILIAN is a statistical wrapper that can be applied to the alignment output file from a spliced aligner and can distinguish false positive junction calls from true positives via assigning a statistical score to each splice junction reported by the aligner.

### 3 File Downloads

Human RefSeq hg38 annotation file was downloaded from: ftp://ftp.ncbi.nlm.nih.gov/refseq/H_sapiens/annotation/GRCh38_latest/refseq_identifiers/GRCh38_latest_genomic.gff.gz

### 4 SpliZ calculation

The SpliZ score computation for a gene consists of two parts: one relative to the 5’ splice sites in the gene (the 5’ splice site SpliZ) and one relative to the 3’ splice sites (the 3’ splice site SpliZ). We first explain how to calculate the 5’ splice site SpliZ for one gene (suppressing the notation specifying the gene for simplicity) and the 3’ splice site SpliZ can be computed similarly. Let *i* specify the 5’ splice site, *j* specify the 3’ splice site, *k* specify the cell, and *ℓ* specify the read. Therefore each junctional read for the gene in the dataset is specified by a unique combination of *ijkℓ*. Note that we only consider junctions for which the 5’ splice site has multiple 3’ splice sites in the dataset.

The 5’ splice site SpliZ score calculation proceeds by treating each 5’ splice site separately. We will consider a plus strand gene and assume that 5’ splice site *i* has multiple 3’ splice sites across the whole dataset (otherwise we would have filtered it out). We rank these 3’ splice sites in order from closest to the farthest from the 5’ splice site in question *i* (i.e. from lowest to the highest genomic coordinate). For example, if there were four 3’ splice sites partnered with *i* across the whole dataset, we would rank them 1, 2, 3, and 4 in the order of their genomic coordinates. For genes that are on the minus strand, the 3’ splice sites are ranked differently as described in the Supplement. Let *r_ijkℓ_* denote the 3’ splice site rank for the read specified by *ijkℓ*. If *r_ijkℓ_* = 1, it means the 3’ splice site has the smallest genomic coordinate among all 3’ splice sites in the dataset.

Now, let *N_i_* be the number of junctional reads observed across all cells for 5’ splice site *i*. We can compute 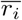, the average rank of the 3’ splice sites for 5’ splice site *i*, as

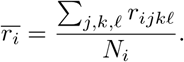

**Figure 1.**
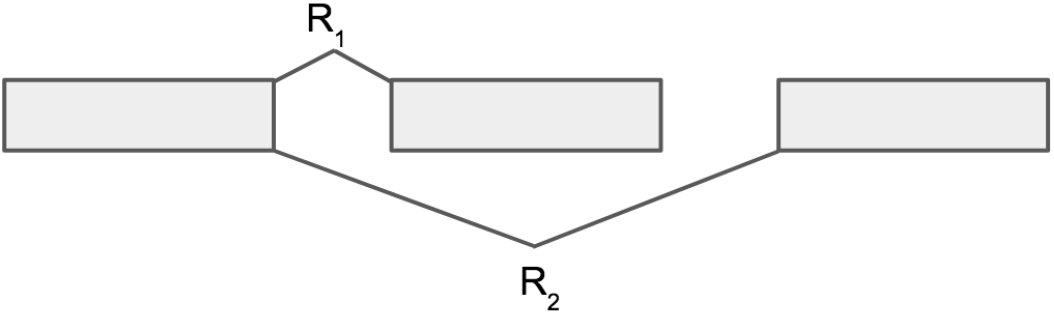
Exon skip with only one junction corresponding to inclusion and one junction corresponding to exclusion

We can also find the variance of the ranks as:

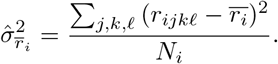

Now, we renormalize the rank *r_ijkℓ_* using the sample variance and mean as:

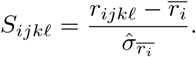

We can see that 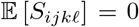 and var (*S_ijkℓ_*) = 1 as we subtract the mean and divide by the standard deviation. The closer the 3’ splice site of a read is to the 5’ splice site, the smaller its corresponding *S_ijkℓ_* value would be. In practice, we truncate the *S_ijkℓ_* values at the 10th and 90th quantiles across all genes to avoid effects from extreme outliers.

We now aggregate these zero-mean and unit-variance variables for all 5’ splice sites in the gene to compute the 5’ splice site SpliZ 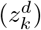 in the *k*th cell as:

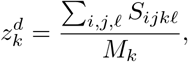

where *M_k_* is the number of junctional reads mapping to the gene in the *k*th cell. It is straightforward to see that 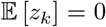. Note that under the alternative hypothesis, for a given cell type 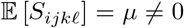. For a cell of this cell type,

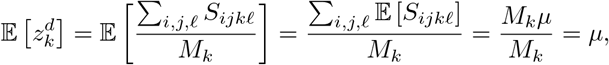

meaning that the SpliZ is not correlated with read depth *M_k_*.

Knowing that the variance of the sum of independent random variables is the sum of their variances:

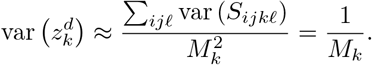

The approximation is due to the fact that the *S_ijkℓ_*’s are not necessarily independent but we expect them to be close enough to independent.

Similarly, we can compute the 3’ splice site SpliZ 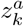. We average the two scores 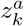 and 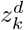 to compute the SpliZ *z_k_* for the gene in cell *k*:

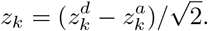

These values are subtracted to correct for signs, such that short introns correspond to small values and long introns correspond to high values for both. Division by 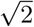 ensures that the variance will be comparable between averaged SpliZ values and SpliZ values for which only one of 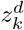 and 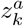 is calculable (in which case that is the SpliZ value for the cell and there is no averaging).

For computing SpliZ, we only consider cells with at least 6 junctional reads mapping to the gene, and the junctional reads for which the 5’ splice site has more than one 3’ splice site observed in the dataset.

### 5 SpliZ PSI equivalence

We will prove the equivalence of the SpliZ and PSI under a specific scenario: when only one of the junctional reads of the exon inclusion event is measured; this could correspond to different ending exons of the transcript (Methods Fig. 1).

Let *n*_1_ be the number of reads for a given cell mapping to the junction between the 5’ splice site and the first exon, and let *n*_2_ be the number of reads mapping between the 5’ splice site and the second exon for that cell.

Then the value of PSI for the cell is 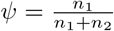.

To calculate the SpliZ *z*, we need to calculate the rank mean and standard deviation across the population of cells.

Assume in the overall population the fraction of junctions including the exon is *f*. Then the mean rank 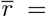 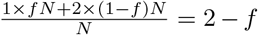 where *N* is the total number of reads mapping to these two junctions across all cells.

The variance *σ*^2^ is given by

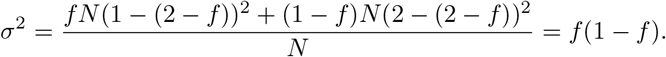

Then the SpliZ value is given by

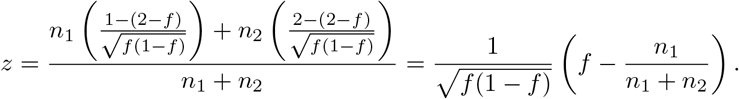

Therefore *z* can be written in terms of *ψ* and *f*:

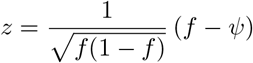

implying that *z* and *ψ* are equivalent in this case.

For example, if *f* = 0.5 then *z* = 1 − 2*ψ*.

### 6 SpliZVD Calculation

Let *M* be an *n* × *p* matrix, where *n* is the number of cells and *p* is the number of splice sites for the given gene. Matrix entries are defined by:

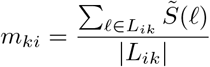

Here *L_ik_* is the set of reads using splice site *i* in cell *k*. 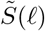 is the normalized residual of the rank of read *ℓ*, defined as follows. For a given gene, let *G* be the set of spliced reads that map to the gene across the data set. Let *S*(*ℓ*) be the residual of read *l*. Then let

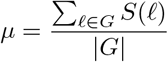

and

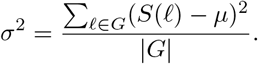

Then 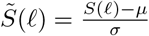, so 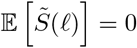 and 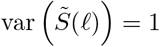.

Let *α*^(*j*)^ be the *j*th eigenvector of *M_ki_*. Then the the *j*th SpliZVD score for cell *k* is given by < *m_k_*, α^(*j*)^ > (note that for this paper we only consider the SpliZVD score based on the first eigenvector, < *m_k_*, *α*^(1)^ >, and refer to this single score as the SpliZVD though more components can be considered). This score has mean 0 and variance 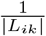 (*j* supressed for simplicity in these calculations):

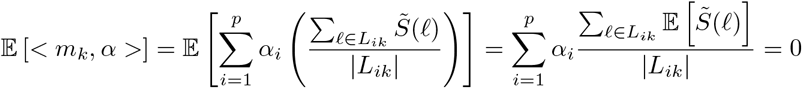

and

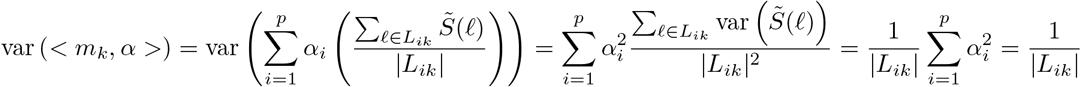

(this is assuming the residuals are uncorrelated).

Under the alternative hypothesis for a cell type, as with the SpliZ, for an individual cell type if 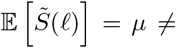 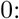

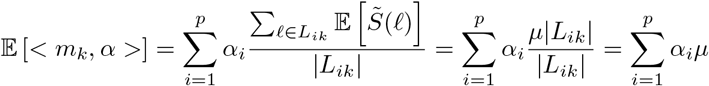

So the SpliZVD value is not dependent on read depth.

### 7 Calling differentially alternatively spliced genes by cell type

We perform the following test independently on each gene for each individual, technology, and score (SpliZ and SpliZVD). The procedure is the same for both SpliZ and SpliZVD, but we will only discuss the SpliZ here for simplicity. For a given gene, we only consider cell types with more than 10 cells with SpliZ values for that gene.

Significance of alternative splicing between cell types in a gene is determined by calculating p values using a two-step procedure based on the work of Chung and Roman [1].

Consider the following equation from Chung and Romano, Lemma 3.1:

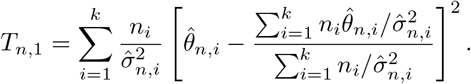

We can compute *T_n,_*_1_ based on the SpliZ values for all cells with splicing expression for the given gene as follows: The *k* samples represent the *k* cell types with splicing expression for the given gene. Then 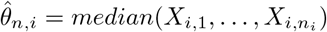 is our test statistic, where 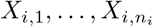 are the SpliZ values for the *n_i_* cells from cell type *i* with splicing expression for the gene. 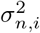 is the sample standard deviation of the SpliZ values for cell type *i*; in practice, we inflate the variance to 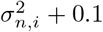 for robustness. Our null hypothesis is that all cell types have the same median SpliZ for this gene.

Performing all permutations of cell types to cells and re-calculating *T_n,_*_1_ yields the permutation distribution (a subset of the permutations yields an approximation of the permutation distribution). Theorem 3.1 states that under some assumptions, this permutation distribution converges to the *χ*^2^ distribution with *k* − 1 degrees of freedom. Also, if the sample distributions do not have different medians, the probability that the permutation test rejects the null hypothesis tends to nominal level *α*.

It is a lot quicker in practice compare to the *χ*^2^ distribution than compute permutations, so we first calculate the p value based on comparing to the *χ*^2^ distribution 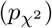. Then if 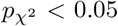 we compute the permutation p value *p*_perm_ by permuting the assignments of cell types to cells for the gene and calculating 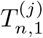 based on this permuted data where *j* is the current permutation. This results in a permutation null distribution 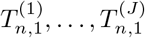 where *J* is the number of permutations performed. Then

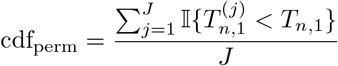

and *p*_perm_ = 2 min(cdf_perm_, 1 − cdf_perm_) (quantifies whether the real value is extreme in either direction). We then adjust the p values for multiple hypothesis testing using the Benjamini-Hochberg procedure. Genes are called as significant if their adjusted p values are less than 0.05.

Genes with the “most deviant SpliZ values” are defined as follows: Find the largest magnitude median SpliZ value *Z_i_* across all cell types for each gene *i*. Then genes with higher *Z_i_* values have more “deviant” SpliZ values.

### 8 Finding the most variable splice sites driving the alternative splicing of a gene

To provide further biological interpretation for the SpliZ and SpliZVD scores and automatically identify the most variable splice sites driving the alternative splicing in a significantly regulated gene, we take the SVD of the residual matrix and use its eigenvalues and eigenvectors to select the most variable splice sites. To do so, we select up to three eigenvectors depending on the values of their corresponding eigenvalues and then in each chosen eigenvector we define up to three splice sites that correspond to up to three elements of the eigenvector with the highest absolute value as the most variable splice site. If the first eigenvalue is greater than 0.7 we select only the first eigenvector and otherwise we select the second and third eigenvectors if their corresponding eigenvalues are at least 0.2. When an eigenvaector is selected we choose the three splice sites corresponding to the top entries that have at least 10% of the eigenvector loadings based on the L^2^ norm of the eigenvector.

Automatic detection of the most variable splice sites in each significant gene re-identified know alternatively spliced splice sites for *CD47* and *TPM2*. Driving splice sites in *CD47* are in chr3-108047292 (a 3’ splice site) and chr3-108057477 (a 5’ splice site), both being involved in annotated alternative splicing events (Figure 3.A). Similarly, the most variable splice sites for *TPM2* are chr9-35684550 (a 3’ splice site) and chr9-35684246 (a 5’ splice site), both known to be annotated alternative exons (Supp. Table *). Also, our SVD-based analysis on *MYL6* identified 56161387 as the most variable splice site, which is involved in a well-known exon skipping event recently shown to be compartment-specific-regulated.

For each driving splice site, we additionally report its annotation status, including whether it is annotated as an alternatively spliced exon. To determine whether an exon is known to be involved in alternative splicing, we extracted the splice sites from the GRCh38 annotation GTF file and then considered those 5’ splice sites (resp. 3’ splice sites) that are observed to be spliced to more than one distinct 3’ splice site (resp. 5’ splice site) across the extracted junctions as known alternatively spliced sites.

### 9 Simulation methods

For each simulation, two cell types are simulated with 20 cells in each. They have pre-defined proportions of each isoform as described in the figure. At each mean read depth *n*, 100 trials are performed, each of which proceeds as follows. First, a read depth for each cell *k* is sampled independently from *Poisson*(*n*). Then for each cell of cell type *c*, the distribution of the *k* reads among splice junctions is drawn from a multinomial distribution, for which each junction from isoform *i* has the probability 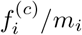, where 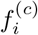 is the underlying fraction of isoform *i* in cell type and *m_i_* is the number of exons in isoform *i*.

Next, the SpliZ and SpliZVD are calculated as described above for this gene and this population of 40 cells. PSI is calculated on a cell-by-cell basis for each cassette exon by dividing the number of reads including that exon over all of the reads that either skip or include the exon in the given cell (note that some reads may not either skip or include the exon; these are disregarded in the PSI calculation).

Then p values are calculated separately for the SpliZ, SpliZVD0, and each PSI value as described above, except *χ*^2^ filtering is not used and variances are not inflated. Benjamini-Hochberg multiple hypothesis testing correction is not used (there is only one score calculated for each of the SpliZ and SpliZVD0; if the multiple PSI values were corrected, it would only cause them to be less significant). The gene is called as having differential alternative splicing between cell types if the calculated p value is < 0.05.

Most variable splice sites are calculated for each by simulating the two cell types at a read depth of 20 and performing the procedure described in section 7.

For each simulation, a “null” simulation is also included, which follows the same setup except both cell types have the same distribution among the isoforms as the first cell type in the original simulation. This allows us to estimate the type I error rate.

### 10 Concordance between datasets

We test whether the number of genes significant in both 10x datasets is more than expected under the null hypothesis, which is that genes found significant in both datasets are due to chance. Let *s* be the number genes that are called as significant in both individual 1 and individual 2 10x data. The probability that a given gene is significant in both datasets is:

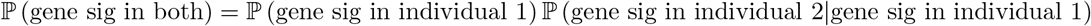

Under the null hypothesis, assume that a gene being significant in one individual is independent from it being significant in the other individual. Therefore under the null hypothesis

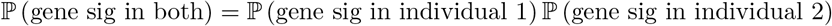

We can estimate the quantities on the right hand side of the question for each individual:

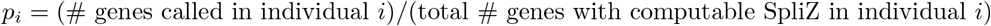

Therefore under the null hypothesis, the probability that ≥ *x* genes are significant in both individuals out of n genes is given by

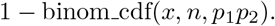

Correlation between datasets is calculated by considering only gene/cell type pairs with at least 10 cells in both datasets, restricting to genes called as significant in both datasets, and calculating the Pearson correlation.

### 11 Cell types “missed” by Smart-seq2

We defined cell types that were almost completely missed through Smart-seq2 sequencing compared to 10x sequencing as SS2-missed cell types. Based on Supplementary Table 2 of the HLCA paper we defined SS2-missed cell types to be those for which

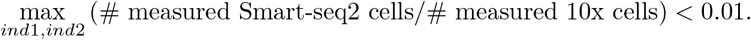

This results in the following cell types as SS2-missed: Proximal Ciliated, Proximal Basal, Proliferating Basal, Mucous, Serous, Capillary Intermediate 2, Bronchial Vessel 1, Bronchial Vessel 2, Lipofibroblast, Mesothelial, CD8+ Memory/Effector T, CD4+ Memory/Effector T, Neutrophil, Mast Cell/Basophil Type 2, Platelet/Megakaryocyt, Macrophage, Proliferating Macrophage, Myeloid Dendritic Type 1, EREG+ Dendritic, TREM2+ Dendritic, Classical Monocyte, OLR1+ Classical Monocyte.

## Notes

https://github.com/juliaolivieri/SZS_pipeline/

### Competing Interest Statement

The authors have declared no competing interest.

### Summary of Updates

Name of the score changed from SZS to SpliZ; modification of the SpliZ, the SpliZVD, introduced; simulations included to compare SpliZ and SpliZVD to PSI; spermatogenesis analysis removed for space reasons; new method of calculating gene-based p values; Figures 1, 2, and 3 revised.

